# Machine learning-assisted identification of factors contributing to the technical variability between bulk and single-cell RNA-seq experiments

**DOI:** 10.1101/2022.01.06.474932

**Authors:** Sofya Lipnitskaya, Yang Shen, Stefan Legewie, Holger Klein, Kolja Becker

## Abstract

**Background:** Recent studies in the area of transcriptomics performed on single-cell and population levels reveal noticeable variability in gene expression measurements provided by different RNA sequencing technologies. Due to increased noise and complexity of single-cell RNA-Seq (scRNA-Seq) data over the bulk experiment, there is a substantial number of variably-expressed genes and so-called dropouts, challenging the subsequent computational analysis and potentially leading to false positive discoveries. In order to investigate factors affecting technical variability between RNA sequencing experiments of different technologies, we performed a systematic assessment of single-cell and bulk RNA-Seq data, which have undergone the same pre-processing and sample preparation procedures.

**Results:** Our analysis indicates that variability between gene expression measurements as well as dropout events are not exclusively caused by biological variability, low expression levels, or random variation. Furthermore, we propose FAVSeq, a machine learning-assisted pipeline for detection of factors contributing to gene expression variability in matched RNA-Seq data provided by two technologies. Based on the analysis of the matched bulk and single-cell dataset, we found the 3’-UTR and transcript lengths as the most relevant effectors of the observed variation between RNA-Seq experiments, while the same factors together with cellular compartments were shown to be associated with dropouts.

**Conclusions:** Here, we investigated the sources of variation in RNA-Seq profiles of matched single-cell and bulk experiments. In addition, we proposed the FAVSeq pipeline for analyzing multimodal RNA sequencing data, which allowed to identify factors affecting quantitative difference in gene expression measurements as well as the presence of dropouts. Hereby, the derived knowledge can be employed further in order to improve the interpretation of RNA-Seq data and identify genes that can be affected by assay-based deviations. Source code is available under the MIT license at https://github.com/slipnitskaya/FAVSeq.

## Background

High-throughput single-cell RNA sequencing (scRNA-Seq) provides a powerful tool for profiling gene expression patterns at single-cell resolution that has revolutionized transcriptomic studies and advanced the knowledge of biological systems. RNA samples for bulk sequencing are typically derived from a heterogeneous population of cells and thus, cell-type specific transcriptomic changes may be lost as the final data represents an average expression across thousands of cells from different types [1]. However, while single-cell technology allows to overcome some limitations of standard bulk RNA-Seq, data complexity and noise increases as well as the detection limit and RNA amount present a challenge, for example, to identify and filter out low-quality genes for reliable and reproducible results. Therefore, in addition to the larger variability of single-cell technology compared to traditional bulk RNA sequencing, the subsequent downstream analysis gives rise to new computational challenges in analyzing data and interpreting the findings.

The increased noise in scRNA-Seq data can be explained by both biological and technical reasons, for example, lower amount of input material, batch effects, amplification biases, cells being in distinct phases of the cell cycle, and transcriptional bursts [2]. Another reason that makes the computational analysis challenging relates to dropout events. Previous studies show that single-cell technology typically suffers from dropouts, i.e., transcripts which exist in a biological replicate or a cell, but are not detected by the sequencing technology. In contrast to RNAs that are simply not present at the time of cell isolation, dropouts have non-zero expression but are not identified due to limitations of experimental protocols as well as other biological and technical reasons [3].

Often these dropouts are caused by low expression, but additional factors may contribute. For instance, dropouts can be potentially caused in different ways by difficulty in isolating single cells from low starting input volumes, cell-specific capture efficiency (e.g., inefficient mRNA to cDNA capture, dilution of cell libraries, and amplification) as well as by the low amounts of mRNA in individual cells [4]. Additional challenges affecting the difference in RNA-Seq experiments can arise from differences in library preparation protocols [5, 6] and computational downstream analysis pipelines [7]. Several studies aimed at understanding the difference between matched RNA-Seq experiments performed a comparative analysis of dataset with available matched bulk and scRNA-Seq samples [8–10]. However, these studies have been are usually limited by the number of sequenced cells and analysed samples [11].

All of that motivated us to perform a detailed comparison of RNA-Seq experiments in order to investigate the most relevant factors affecting the difference between them. To do so, we analysed paired data—scRNA-Seq and bulk from the same sample—to limit the biological sample-to-sample variation. We focus on the analysis of the difference among gene expression measurements and also provide FAVSeq, a machine learning (ML)-based pipeline, which identifies features (e.g., genomic factors) affecting variability in matched single-cell and bulk RNA-Seq data. Results suggest that 3’-UTR and transcript lengths influence the gene expression difference between experiments at the most. Likewise, these features together with cellular compartments were found to be relevant for global dropouts. Thus, the analysis presented in this paper provides the basis for further experimental investigations of identified factors, as well as the subsequent improvements at the level of RNA-Seq experimental analysis and data pre-processing that allow to facilitate the fundamental research and biomedical applications based on RNA sequencing technologies.

## Results

### Matched bulk and single-cell experiments allow for a detailed analysis of differences in gene expression measurements

With the aim to assess sources of variation provided by different RNA-Seq technologies, we based our analysis of gene expression measurements performed in multiple biological replicates (hereinafter referred to as samples). Thus, considering distinct samples allows for improving the efficiency of statistical testing and the reliability of the findings. In order to perform a comparative analysis of bulk and scRNA-Seq experiments, both sequencing technologies were applied to eight retina samples (6 domestic pigs and 2 minipigs). The unified sample preparation procedures ensured high comparability of resulting measurements. Specifically, all samples were processed according to the (10x Chromium) single cell RNA-Seq sample preparation protocol—including single cell dissociation—before fractions were split for the two experiments providing matched scRNA-Seq and bulk RNA-Seq measurements. Subsequently, RNA-Seq libraries for bulk sequencing were prepared using the NEB-Next Ultra mRNA, followed by paired-end Illumina sequencing (2 × 75 bp). The scRNA-Seq experiment, performed using paired-end sequencing 10x Single Cell 3’ v2, produced a total of 2,111,208 cells with an average number of 263,901 cells per sample (Supplementary Figure S1A). For a more detailed comparison of experimental protocols of the matched RNA sequencing experiments see Methods and Supplementary Table S1.

In order to compare gene expression measurements provided by different experiments, we aggregated scRNA-Seq data by summing up raw read counts across single cells from each sample, followed by its gene length and per sample normalization, which we termed “pseudobulk”. The latter was shown to be highly similar (*ρ* = 0.96) to original measurements provided at the single-cell level (Figure S1B). Out of 25,322 annotated genes in the Sus Scrofa genome, 21,372 genes were detected and showed non-zero expression in at least one out of eight samples in either of the matched experiments. Among all samples, 19,965 and 19,436 genes were detected in at least one sample of the scRNA-Seq and the bulk experiment, respectively (Figure 1A). Specifically, in the bulk experiment we found an average number of 17,283 genes per sample, while the scRNA-Seq provided the average number of 16,733 (Figure 1B), indicating a slightly lower detection rate for the single-cell protocol. Among these detected genes, 18,029 (*>* 71 %) were common, i.e., showed non-zero expression in at least one sample of both scRNA-Seq and bulk measurements (blue intersection in Figure 1A). At the same time, we observe genes with missing expression in one experimental modality (e.g., scRNA-Seq), while being detected in the matched sample provided by another experiment (e.g., bulk RNA-Seq), which we term as sample-wise dropouts (gray areas in Figure 1A).

**Figure 1.**
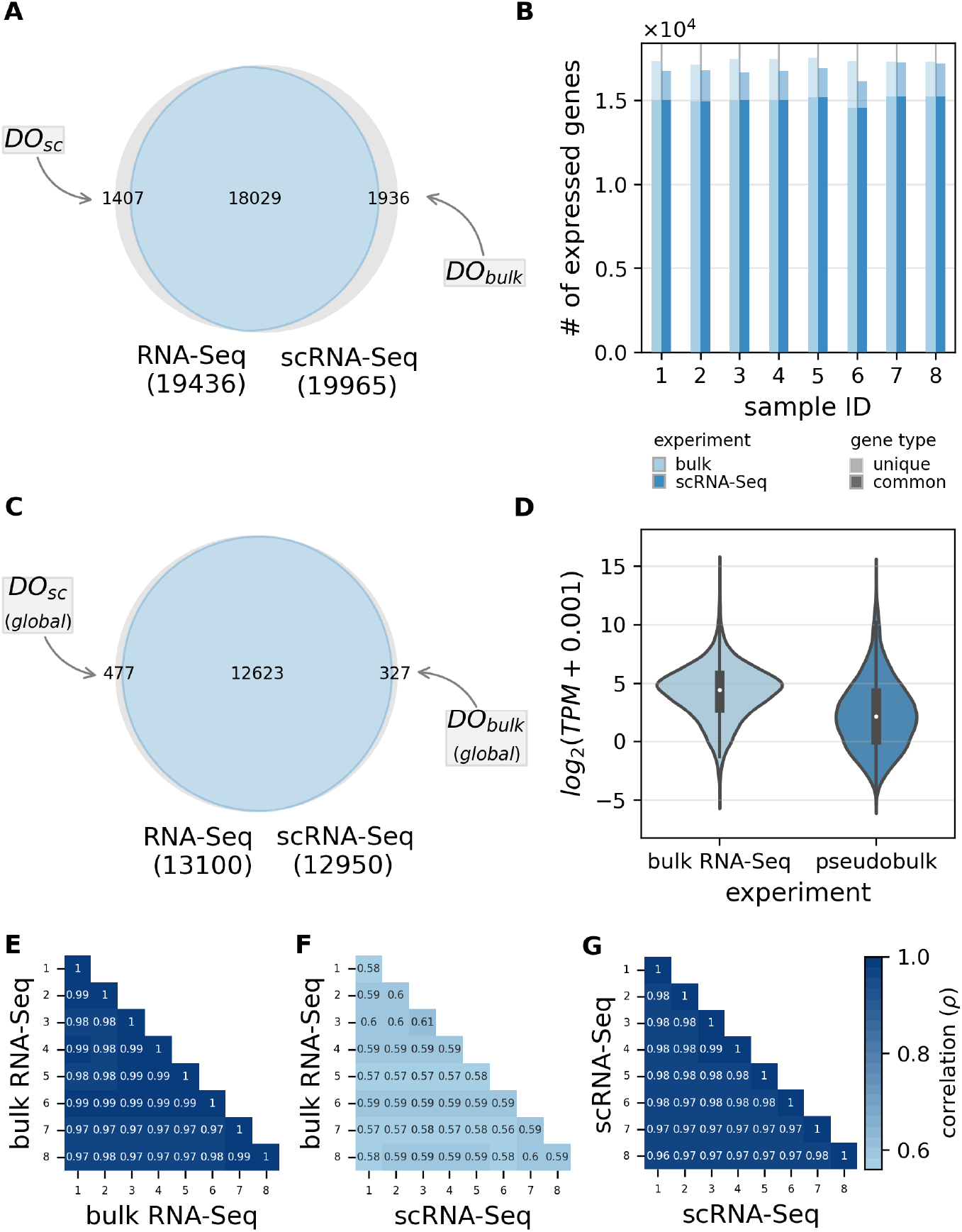
Comparison of the bulk and single-cell gene expression profiles in matched RNA-Seq experiments. A. Amount of uniquely expressed genes detected in the bulk and scRNA-Seq, where blue color represents genes which are expressed in both experiments and gray indicates dropouts (DO) or genes with missing expression across all samples in the one of experiments. B. Quantitative analysis of common and unique (missing in one of the experiments) genes detected by different RNA-Seq technologies. C. Amount of genes detected either in all samples in both RNA-Seq experiments (intersection area, blue), or missing genes that are expressed in 8 samples (a.k.a. global dropouts) in only one of the matched experiments (disjunctive union, gray). D. Distribution of averaged expression measurements of most confident common genes detected in all samples (blue intersection in Figure 1C) in the bulk and aggregated scRNA-Seq experiments. E. Sample-wise correlation of transcriptomic profiles of common genes (blue intersection in Figure 1C) measured in the bulk experiment. F. Sample-wise correlation of transcriptomic profiles of common genes measured in the matched dataset. G. Sample-wise correlation of transcriptomic profiles of common genes measured by scRNA-Seq.

Analysis of gene sets in individual experiments indicates (Figure 1C) that the bulk RNA-Seq provided a slightly higher proportion of the most confident genes that were detected across all samples, i.e. 97.6 % (13,100) of genes were detected by the bulk RNA-Seq in all samples against 96.4 % (12,950) detected by the scRNA-Seq experiment. The total of 12,623 genes are found to be common or expressed in all available 16 samples across matched experiments (blue intersection in Figure 1C). Besides, we also found genes with the available expression values in 8 samples in only one of the experiments (referred later as global dropouts). Accordingly, both singlecell as well as bulk global dropouts exist with the total of 804 genes (gray areas in Figure 1C) that have been systematically found in the combined experiments (Figure S1C).

We further found a profound difference in expression profiles between bulk and pseudobulk measurements. While many common genes were detected in matched experiments, the quantitative expression levels differ between bulk and pseudobulk datasets at the global level. Comparison of mean expression levels of genes detected in both matching experiments (Figure 1D) indicates the higher detection rate in the bulk experiment demonstrating the consistency with reported studies [3]. Figure 1D shows a substantial difference in expression measurements with lower gene expression measured in the single cell experiment with an average of *log*_2_ *TPM* = 2.14 when compared to the bulk experiment with median expression of *log*_2_ *TPM* = 4.44.

As the ground-truth of expression in genes with missing values is unknown, we can only obtain such an information from samples of the matching experiments. Consequently, in order to ensure the validity of the derived results, we performed an ML-assisted analysis using the subset of the most confident genes (Figure 1C). Thus, for the analysis of the quantitative difference we used data related to common genes with available expression measurements in all samples in both RNA-Seq experiments (blue intersection area; regression task), while for the analysis of the qualitative difference (dropouts) we also include those with expression measurements detected in 8 samples in the matched experiments only (gray areas; classification task).

### Technical noise is the major contributor to difference between single-cell and bulk RNA-Seq

In order to assess the contribution of technical and biological variation, we performed a sample-wise correlation analysis within and between the two sequencing experiments. Figure 1E-G indicates how transcriptomic profiles produced from the scRNA-Seq experiment are different to that of the bulk RNA-Seq for genes expressed across all samples and both modalities (blue intersection in Figure 1C). For each of the sequencing protocols, the expression measurements showed a little across-sample variation with correlation coefficients close to 1 in both the bulk RNA-Seq (*ρ >* 0.97) and aggregated into pseudobulk scRNA-Seq (*ρ >* 0.96) experiments as represented in Figure 1E and Figure 1G, respectively.

This suggests that the effect of the biological noise is negligible in the tested setup. Results also indicate a slightly increased gene expression variability level in aggregated single cells compared to bulk RNA-Seq. As droplet based scRNA-Seq systems (i.e., 10x Genomics Chromium) amplifies cDNA fragments close to polyadenylation (polyA) tails, the corresponding gene expression measurements are highly biased to the 3’-end, while the full transcript coverage was captured in the bulk data. Due to the generated single-cell data are often confounded by the quality of 3’-UTR annotation, this increase in variability of gene expression measurements was expected.

On the other hand, correlation between bulk and pseudobulk samples was comparatively low (*ρ* = 0.59 ± 0.01), suggesting the presence of technical variation between matched RNA-Seq experiments (Figure 1F). Thus, the correlation analysis reveals a negligible impact of batch effects on gene expression indicating that technical noise is the major contributor to the difference between matched RNA-Seq experiments.

### Dropouts are systematically observed in data and are only partially caused by lowly expressed genes

Dropout events relate to a common phenomenon observed in RNA-Seq data implying that specific transcripts cannot be detected by the sequencing technology [4]. As the presence of dropouts highlights possible limitations in the sequencing and/or pre-processing protocols, which, in turn, may introduce bias in downstream analysis and interpretation of the data, we investigated these genes in more detail.

Supplementary Figure S1C provides an overview of the experiment-wise sparsity rates of dropouts identified in the matched dataset, where the sparsity rate is defined as the number of genes with missing expression (*TPM* = 0) across different proportions of samples. Results of this analysis together with Figure 1B indicate that sample-wise dropouts are not likely to occur randomly or by chance in matched RNA sequencing data. Here, the sparsity rate of zero indicates dropouts with the available expression values in all 8 samples in only one of the experiments.

Given that our experimental design provided gene expression measurements across several biological samples, we introduce a more specific definition of high-confidence dropouts (here and later as global dropouts), in which we consider them as genes representing missing expression in all samples in only one of the matched RNA-Seq experiments, while being detected (*TPM* ≠ 0) across all samples by another technology. Based on eight matched bulk and pseudobulk samples in our dataset, we found 804 global dropouts, with a higher number in the single cell experiment (gray areas in Figure 1C and bar 0 in Figure S1C). Specifically, a quantitative analysis of global dropouts indicates 477 genes were not detected in scRNA-Seq data, while being expressed in all samples of the bulk data. Conversely, 327 genes show no expression in any of the bulk samples, while being strongly expressed across all samples in single cells.

Since the previous research suggests that weakly expressed genes tend to produce more differences than highly expressed genes at the single-cell level [2], we analyzed mRNA expression level of global dropouts in the matched experiment (Figure S1D). Subsequently, comparison of expression measurements indicates the similar average expression levels of these genes in the bulk and scRNA-Seq experiments. As expected, we also observed the majority of global dropout genes to be lowly expressed in the matched experiment. However, a substantial number of dropouts showed non-marginal (high) expression, as can also be seen for specific examples. Closer examination of global dropouts that are highly expressed in the matched experiment (Figure 2A and Figure 2B for single-cell and bulk RNA-Seq data, respectively), reveals that these genes represent variable expression patterns and therefore, cannot be explained by the low expression level only.

**Figure 2.**
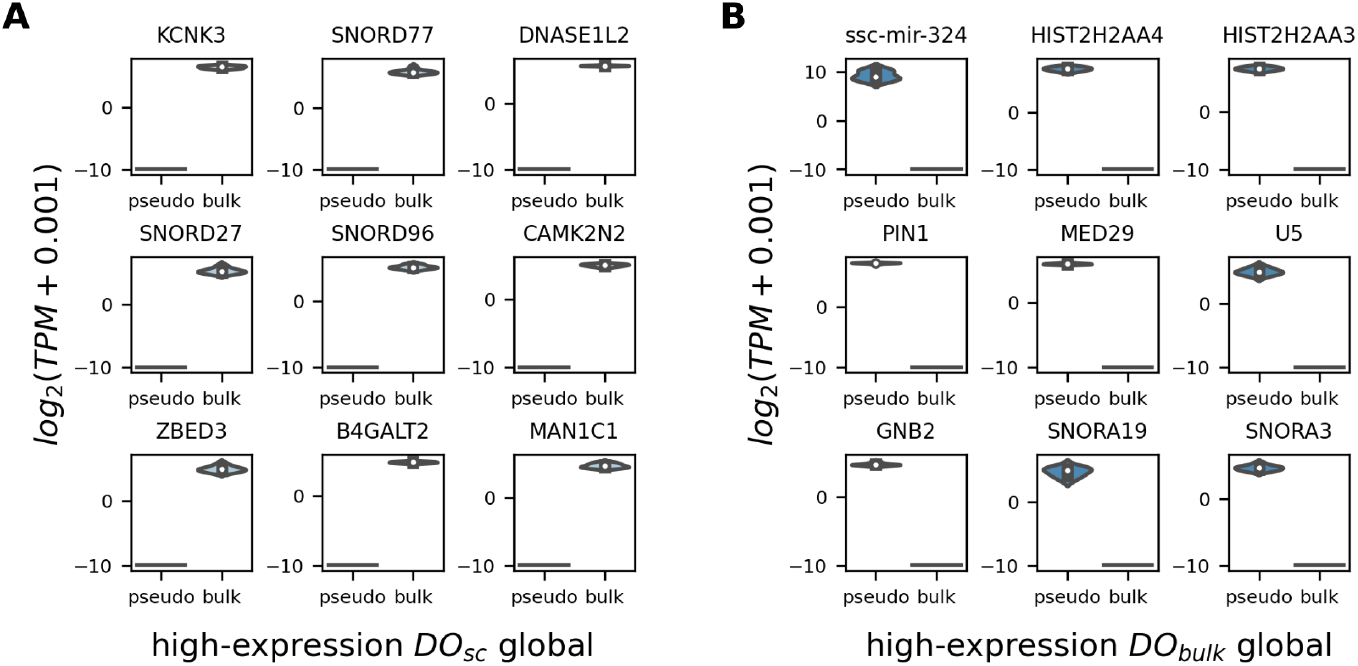
Gene expression levels of most confident global dropouts. Measurements in the matched experiment are shown. A. Expression levels of genes that are global dropouts in scRNA-Seq and exhibit a high-expression in the bulk experiment. B. Expression levels of genes that are global dropouts in the bulk experiment and exhibit a high-expression in scRNA-Seq.

### Proposing a computational approach to analyse the difference in RNA-Seq experiments

In order to identify which factors determine whether genes are differently detected in matched RNA-Seq experiments, we introduce FAVSeq (Factors Affecting Variability in Sequencing data), a machine learning-assisted pipeline, those design intends to support researchers in disclosing potential root causes of the difference—in terms of gene expression measurements and dropouts–observed between RNA-Seq technologies. FAVSeq enables to select features obtaining the strongest predictive power for estimation of technical variability between RNA sequencing modalities. The pipeline includes the following steps (Figure 3):

**Figure 3.**
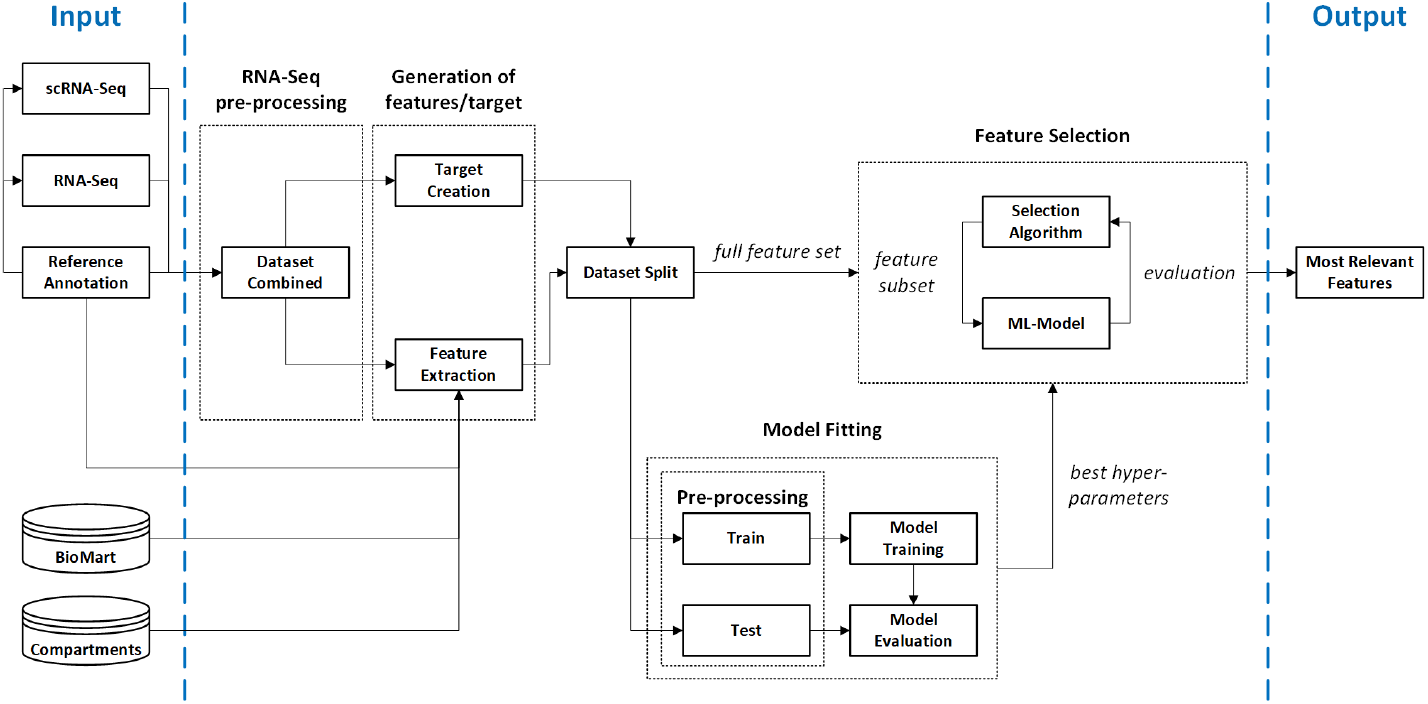
Framework utilized in the FAVSeq pipeline for ranking and selection of features affecting the technical variability in RNA-Seq datasets of matched experiments. The raw input acquired from different sources (on the left) is being pre-processed in order to build a set of features to be fed into an ML-model (e.g., random forest). The full set of input samples is used to perform search for optimal hyper-parameters of the ML-model using 5-fold CV. The chosen hyper-parameters serve then to perform selection of most relevant features based on the underlying ML model (framework’s output, on the right) using RFE feature selection technique that allows to determine most informative features, according to the ML model’s scores.

1. Create the target by calculating the ordinary least squares (OLS) residuals in gene expression levels.
2. Generate gene-associated features based on GTF annotation file and openaccess databases (e.g., BioMart, Jensen Compartments).
3. Optionally, recover missing values in features using a chosen imputation approach (e.g., model-based one).
4. Optimization of hyper-parameters of models through the 5-fold cross-validated (CV) grid-search. Selection of the top-performance model.
5. Rank features w.r.t. their influence on the objective using the top-performance model and based on recursive feature elimination (RFE) in 5-fold CV.
6. Output the summary reports, including statistics for the core features that contribute to the difference between experimental modalities.

Thus, steps 1-3 provide us with the independent (features) and dependent (quantitative or dropouts-related target differences) variables. Feature selection is not a trivial task, especially in case of genomics databases that often consist of partially incomplete data, which, in turn, may affect the performance of machine learning models. To address this issue, we integrated a missing data imputation module into FAVSeq (step 3). The introduced module supports both non- and parametric imputation strategies, including k-Nearest Neighbors (kNN), that was shown to be effective to handle missing and/or corrupted values in genomic and transcriptomic data [12–14].

Then, based on the created dataset, the subset of the most relevant features is being selected using the machine learning-assisted feature selection approach (steps 4-5), which, in turn, consists of two main parts. The first part serves to select the most suitable model to predict the target difference between experiments (training, evaluation and optimization of hyper-parameters), and the latter part serves to select the subset of the most relevant features among the set of tested ones w.r.t. the objective (Mean Squared Error or Balanced Accuracy for regression and classification tasks, respectively).

While analysing features w.r.t. quantitative difference (regression task), the random forest model was used because it’s agnostic to variable types (numerical or categorical). Furthermore, given that smallest and largest values of some features differ by several orders of magnitude, such a model can provide sufficient number of estimators, so individual trees cover particular ranges of the input. Also, the model preserves monotony of transformations applied to the input variables, and is not sensitive to outliers in the input samples. For the classification-based analysis of dropouts-associated features, we used a multilayer perceptron (MLP) model, those training and evaluation followed the same steps. To derive an optimal subset of features relevant for the difference between experiments, we utilized RFE, which is a model-agnostic meta-learning method allowing to exclude the least important ones according to the model’s scores on each iteration. The algorithm works until the acquisition of the most important features w.r.t. the objective (target difference).

Finally, the pipeline provides the summary reports (step 6) in a form of tables and visuals (i.e., best hyper-parameters chosen, comparison of the performance of models), which allow interpreting the obtained results as well as guiding on further experiments to be performed.

### Aggregating gene expression difference in matched RNA-Seq experiments

In order to define a quantitative metric for comparing RNA-Seq sequencing experiments, the aggregated per-sample gene expression difference was calculated and used as a dependent variable for the regression analysis. Specifically, we calculated the averaged minimized sum of square differences between measured gene expressions in bulk and aggregated single cells using the OLS method, as described in Methods section. Log-transformation of expression values was done beforehand to make them conform more closely to the normal distribution.

Figure S2A represents how the quantitative difference—residuals of gene expression measurements—is estimated for the sample one for genes found in pseudo- (y-axis) and bulk (x-axis) RNA-Seq datasets, showing the higher measurements in the bulk accordingly with the analysis depicted in Figure 1D. Gray color indicates dropouts and blue indicates genes detected in both matched RNA-Seq experiments for the total of 13,427 genes. At a global level (Figure S2B), we further notice that the quantitative difference—averaged residuals of gene expression measurements— aggregated across all samples is also higher for genes in the bulk experiments in comparison with those in the aggregated single cells. Thus, the calculated quantitative difference indicates how close measurements in matched experiments are by relating gene expression in pseudobulk to bulk RNA-Seq and serves as a basis for further regression-based feature selection analysis. At the same time, the dropouts-related difference (non-/dropout) will be used later for binary classification task while identifying features contributing to the presence of dropout events.

### Creating the feature representation of genes by aggregating data from genomic databases

In order to identify the most relevant features for the difference between gene expression measurements, we generate prospective features representing factors for genes available in both experiments (listed in Table 1). These features comprise the subcellular localization of a gene product, the chromosome at which the gene is located as well as metrics describing the dimensions (e.g., transcript length, UTR length) or expression of a gene (transcript count). Collection of data from genomic databases for the further feature engineering was done for 12,623 common genes that are expressed in all samples across matched experiments (blue intersection in Figure 1C), also considering 804 global dropouts (gray areas in Figure 1C) for the further identification of features affecting the quantitative difference and dropouts events, respectively.

**Table 1.** Summary of the generated gene-specific features

| Feature name | Description | Sources |
| --- | --- | --- |
| **compartment** | subcellular localization | COMPARTMENTS [15] |
| **TSS** | transcription start site | BioMart [16] |
| **GC-content** | guanine-cytosine content ratio | BioMart [16] |
| **cDNA length** | length of complementary DNA | BioMart [16] |
| **transcript count** | number of transcripts | BioMart [16] |
| **chromosome** | chromosomes/scaffolds id | GTF (Ensembl v.86) |
| **transcript length** | length of transcript | GTF (Ensembl v.86) |
| **3'UTR length** | length of 3'untranslated region | GTF (Ensembl v.86) |
| **5'UTR length** | length of 5'untranslated region | GTF (Ensembl v.86) |
| **CDS length** | length of coding DNA region | GTF (Ensembl v.86) |
| **biotype** | transcript variant type (ex. coding) | GTF (Ensembl v.86) |

For this purpose, a diverse data corresponding to genes were integrated from different sources, such as COMPARTMENTS [15], BioMart [16] and GTF (Ensembl v.86) annotation data [17]. Since the chosen features were of both numerical and categorical types, the latter ones were transformed into numerical representation by enumerating the categories and assigning the corresponding indices. As a result, we created a set of 10 prospective features representative for matched sequencing experiments.

Since machine learning models greatly benefit from high quality training data, it is necessary to assure such a property of the input data. One of the factors that can be potentially harmful to ML-models is multicollinearity, i.e. statistically non-independent relationships between features. In order to determine whether additional feature selection is required as a pre-processing step before the use of the FAVSeq pipeline, we performed a cross-correlation analysis that allowed to assess the occurrence of high inter-correlations among independent variables. Its results suggest (Supplementary Figure S2C) that filtering of the features is not necessary, since they do not exhibit strong dependencies between each other [18]. Additionally, we performed the comparison of pairwise feature correlations according to Pearson and Spearman (upper rights and lower left in Figure S2C, respectively) that suggests the existence of non-linear relationships between some features (e.g., 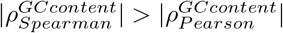), which we consider later while choosing among suitable machine learning models. With the aim to disclose a subset of the most relevant features for the aggregated target difference, we performed an ML-assisted analysis.

### Identification of factors affecting the gene expression difference in RNA-Seq experiments

Using the proposed FAVSeq, we analysed the influence of generated features on bulk vs. scRNA-Seq gene expression differences calculated for the matched experiments. For that, we considered 5134 genes with no missing values in any tested features (data of 0% sparsity; Table S2). The model performance on the gene expression difference prediction task was assessed using MSE that served as a loss metric. Additionally, we used simple linear regression as a baseline in order to show how the errors are measured on the same data in comparison with those calculated based on analysis using random forest with optimized hyper-parameters. Comparison of the performance of regression models indicate the random forest as the most suitable model to predict the quantitative difference between experiments based on the set of tested features (Figure 4A). Here, the best performance was achieved using an ensemble of 512 trees with the depth of 8. During the model training, at least 16 samples were needed before splitting tree’s internal nodes, while leaf nodes contained 4 or more samples.

**Figure 4.**
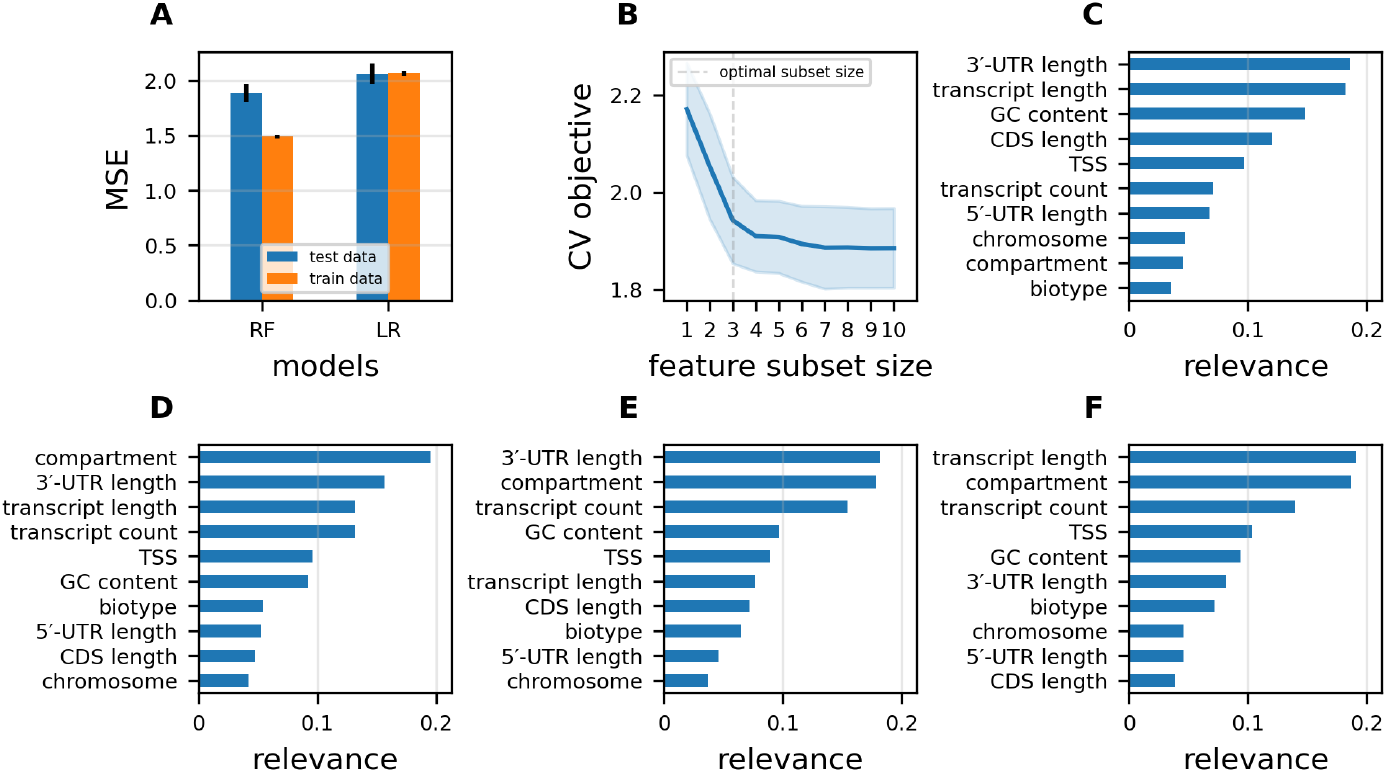
Identification of features contributing to the quantitative and qualitative (dropouts) differences in the matched RNA-Seq experiments. A. Comparison of the performance of machine learning models—random forest (RF) and linear regression (LR)—in prediction of the quantitative difference based on tested features in terms of MSE (lower is better) in 5-fold CV. B. Solution on the optimal number of features based on RFE-RF used to examine feature importance across different subset sizes (x-axis). Cross-validation loss (y-axis) indicates how the model performs based on differently-sized subsets of features in 5-fold CV, showing the steep drop in mean loss values around the subset of three features. Dashed vertical line indicates the optimal number of features. C. Ranking results for features affecting the quantitative difference between experiments. X-axis indicates feature relevance values derived using the RFE-based-on-RF model and y-axis indicates feature names. D. Ranking results for features affecting dropouts in both experiments and derived using RFE-MLP. E. Ranking results for features affecting dropouts in scRNA-Seq and derived using RFE-MLP. F. Ranking results for features affecting dropouts in the bulk RNA-Seq and derived using RFE-MLP.

Subsequently, we derived an optimal subset of relevant features using recursive feature elimination based on random forest (RFE-RF) to obtain rankings within differently sized subsets of features in a 5-fold cross-validation. Figure 4B indicates how well the difference can be explained by the tested subset of features based on the RFE-RF approach. The cross-validation loss curve (lower is better) shows that the model error decreases with the number of features used and reaches the global minimum when using a subset of size 9. Then we determined the optimal number of features through the search for the first stationary point of the objective function by calculating second-order differences, which indicate the steepest drop of the curve. We see noticeable deceleration of the drop of the loss curve after the feature set’s size reaches value of 3, indicating these features as the most important ones w.r.t. the difference discovered between the RNA-Seq experiments (dashed vertical line in Figure 4B).

The same model was then used to rank features w.r.t. their influence on changes in the target variable. As random forest ranking considers both feature set completeness and non-linear interactions within it, we drawn the conclusions about the feature importance upon its scores. Closer look at the importance scores indicates the major impact of three particular features—3’-UTR, transcript length and GC content—on the quantitative difference between matched RNA-Seq experiments, as also shown in Figure 4C. In total, the aforementioned factors are responsible for more than 51 % of the entire relevance of the features for the regression target.

In order to provide a more in-depth understanding on how these features may vary between subsets of genes, we calculated first and second central moments of the top-identified features for the least and most different genes according to the previously calculated measure of the difference in the matched experiments, as shown in Table 2. Interestingly, we observe the shortened length of 3’-UTRs in the most different genes, suggesting the higher level of gene expression as well as the possible differences in their mRNA metabolisms (e.g., sub-cellular localization, stability and the rate of translation of mRNAs [19, 20]).

**Table 2.**
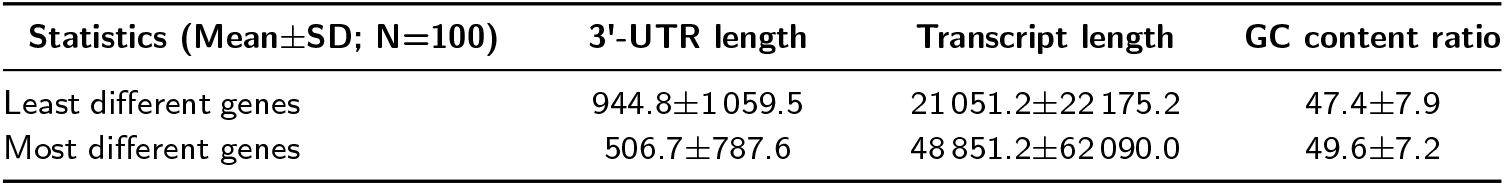
Characteristics associated with top-relevant identified features in least and most different genes according to the analysis of matched RNA-Seq experiments

### Identification of factors affecting the presence of dropouts in RNA-Seq experiments

In the previous Sections, we focused on quantitative expression differences between single cells and bulk RNA-sequencing. Here, we applied the similar approach introduced in the FAVSeq pipeline for the classification task in order to identify the most relevant factors associated with the occurrence of dropouts. We asked whether the generated features in Table 1 affect the presence of dropouts (qualitative difference) in RNA-Seq experiments and therefore we adapted our machine learning approach to this group of genes. For this analysis we chose 13,427 genes, including 804 global dropouts, representing expression measurements across all samples in the opposite experiment (Figure 1C, gray areas). As the features generated in the previous step consist of missing values (Table S2), these values were imputed using a k-NN model as it was shown to be more accurate on dropouts prediction task (Figure S2D).

Unlike regression, the classification task implies estimation of discrete (class labels) rather than continuous values. In a such setup, one has to take into account class imbalance, since strong domination of one class may introduce severe bias into the model’s predictions. As the presence of a dropout—and the corresponding class in the data—is a quite rare event (*<* 6 %), the complementary class, namely the absence of a dropout, is highly over-represented, since the corresponding samples occur roughly 15.6 times more frequently (Table S2). To address this issue, one can utilize the oversampling approach for the training data or choose a model, those training procedure allows to re-weight classes assigning higher importance to the under-represented class (dropouts). The latter is possible using an MLP model trained using weighted cross-entropy loss as the optimization objective. In order to account the class imbalance during the performance evaluation, we also applied class-wise averaged recall (a.k.a. balanced accuracy) to assess the model’s prediction accuracy. Results of the feature ranking w.r.t. global dropouts found in both experiments indicate that cellular compartments, 3’-UTR and transcript lengths affect the presence of dropouts in the combined bulk and scRNA-Seq dataset (Figure 4D). The best model performance was achieved for the MLP containing 512 hidden neurons trained using learning rate *η* = 0.01 and regularization term *α* = 0.0001. Exact values of the aforementioned hyper-parameters were obtained through the grid search over a predefined hyper-parameter space. Furthermore, we utilized this approach to analyse features w.r.t. their influence on experiment-wise global dropouts (including 477 and 327 in scRNA-Seq and bulk, respectively). Latter results indicate the similar factors to be relevant for dropouts in individual single-cell (Figure 4E) and bulk (Figure 4F) RNA-Seq, while the 3’-UTR exhibits a particular importance for dropouts in the single cell experiment.

## Discussion

The proposed here FAVSeq pipeline allowed to investigate the impact of a variety of factors (i.e., genomic and transcriptomic) on the variability in RNA-Seq data of matched experiments. Our results suggest that 3’-UTR and transcript lengths, as well as GC content influence the gene expression difference between the bulk and scRNA-Seq profiles. Similarly, 3’-UTR and cellular compartments were found to be relevant for dropouts at the most. Some of these identified features have been already reported to affect gene expression variability in RNA-Seq data. In particular, 3’-UTR was shown in determining expression differences in RNA-Seq.

From the technological standpoint, the potential reason for 3’-UTR to be appeared as one of the core factors affecting gene expression variability is that reads can be biased towards the 3’-UTR in single cell sequencing. Droplet-based single-cell RNA-Seq methods (e.g., 10x Genomics Chromium), have the majority of the reads mapped to UTR (mainly 3’-UTR) [21], while in the bulk both ends are typically used for tagging of mRNA fragments (cell-barcode + UMI). This could lead to the result we see, that 3’-UTR length contributes to a difference between the expression levels. Thus, the 3’-UTR in such a case should be more relevant in measuring gene abundance. Moreover, in the case of the pig data the 3’-UTR can be identified as the most influential feature because this non-model organism has less well annotated 3’-UTR regions [22], while bulk doesn’t have 3’-bias, so it wasn’t affected so much. In order to explore deeper the differences between scRNA-Seq and bulk, one can further utilize FAVSeq for the matched data from non-model organism (e.g., mouse as provided by Chen et al., 2020 [23]).

Additionally, the following approaches can be employed to extend the current analysis at different levels (data preparation and analysis steps), such as: A) test how additional features can affect the gene expression variability; B) further optimize the hyper-parameters of the model; C) try out other algorithms. With regards to the data preparation step, additional features extracted from the unprocessed sequencing data (e.g., BAM, FASTQ) can be considered. Thus, the feature selection analysis based on data with the least of downstream processing steps will allow to investigate in more detail the experiment-related factors affecting the technical variability between sequencing protocols. Another possibility addressing individual model’s performance in the data analysis step is to further explore hyper-parameters of the model during the optimization phase. Thus, the performance of the model can be further improved by searching for more optimal values in the increased hyper-parameters’ space. Accordingly, the number of hyper-parameters can be expanded and/or the model can be calibrated by the analysis of out-of-bag (OOB) errors that serves as cross-validation loss to approximate a suitable number of trees at which the error stabilizes.

Furthermore, the ML pipeline can be extended by testing other machine learning algorithms and meta-learning approaches (e.g., Support Vector Machine, Boosting, k-Best) to achieve higher predictive power. For instance, boosting of the regression trees can be used in order to improve further the model performance on the gene expression difference prediction task. Thus, the FAVSeq-based analysis can be extended by, for instance, introduction of additional features (e.g., cell specificity)— that are dependent on the scientific question formulated by a researcher—in order to verify their relevance for the difference between RNA-Seq technologies of interest.

## Conclusions

In this study we demonstrated computational framework to investigate the sources of variation in RNA-Seq profiles obtained from the same population of biological replicates based on different RNA sequencing technologies. Based on the analysis of matched single-cell and bulk RNA-Seq data, we found high similarity in gene expression measurements for the majority of genes and also discovered that this difference is primarily subjected to a technical variation given the negligible effect of biological variation in the tested setup. In addition, we performed an in-depth analysis of dropouts which were found to be systematically present in both experiments and to be not explained by low-expression genes only, as it was generally accepted in the preceding studies.

Furthermore, we proposed an ML-based pipeline, namely FAVSeq, for detection of important factors (i.e., genomic and transcriptomic) affecting quantitative (gene expression levels) and qualitative (dropouts) difference in matched RNA-Seq experiments. We analysed matched single-cell and bulk dataset to discover that 3’-UTR and transcript lengths affect gene expression variability between these sequencing experiments at the most. Subsequently, we applied FAVSeq for identification of features associated with the occurrence of dropouts and found out the same features together with cellular compartments to be relevant for presence of global dropouts as well. We also demonstrated how to reconstruct missing values in data generated from metadata and genomic databases based on the k-NN approach.

Therefore, the proposed computational framework enables to investigate the sources of variability in RNA-Seq experiments, which, in turn, allows to improve the interpretability and reduce the complexity of the further in-depth analysis of gene expression data provided by different RNA-Seq technologies.

## Methods

### Data sets

To analyse the core factor affecting the difference between RNA sequencing experiments, we examined single-cell (scRNA-Seq) and mRNA (bulk RNA-Seq) datasets measured on the same population of retina cells extracted from eight pig animals (Sus scrofa) and provided in the study of Shen et al., 2021 [22].

#### Library preparation and processing of RNA-Seq datasets

In short, single cell suspension from retina was originated from 8 pig animals (6 pigs and 2 mini pigs), followed by library preparation done using the Single Cell 3’Reagent Kit v2 (10x Genomics). Matched sequencing experiments were performed on the same biological samples to measure gene expression patterns. In order to mitigate the difference between sequencing procedures, the same scRNA-Seq cell preparation procedure was used in both experiments. For each library pool the sequencing was performed on the Illumina HiSeq 4000 platform to generate scRNA-Seq and the bulk dataset with single-indexed paired-end and dual-indexed 2×75 bp paired-end runs, respectively. Sequencing reads for both datasets were mapped and annotated on a gene level using the same version of Sus scrofa reference genome (version 10.2.86 primary assembly). Transcripts per million (TPM) was calculated subsequently to measure gene abundance in the RNA-Seq experiment.

### Experimental setup

In order to identify the most relevant contributors to the variability in RNA-Seq experiments, we utilized embedded and wrapper feature selection methods to access the importance of a variety of factors (i.e., genomic and transcriptomic) on the quantitative (gene expression levels) and qualitative (dropout events) difference in matched bulk and single-cell RNA-Seq dataset.

#### Creation of pseudobulk from scRNA-Seq

In order to enable the comparative analysis of data from different experiments, single cell counts were aggregated by summing up raw read counts across single cells from each sample, followed by its gene length and per sample normalization giving transcript counts per million.

#### Extraction and aggregation of gene-associated features

Data corresponding to genes were integrated from different sources, such as COMPARTMENTS [15], BioMart [16] (v.86) and Ensembl v.86 (parsed from GTF) annotation data [17]. As features representing a length (i.e., transcript length) consist of duplicates, the longest entries were chosen for the analysis. For the same reason (presence of duplicates), an averaged GC content was calculated for the corresponding feature. Chromosome feature provides the information on chromosome localization, where numerical entries (i.e., chromosome 2) are designated by their chromosome numbers.

A feature representing cellular compartments was generated as described below. First, genes with the most reliable entries were selected w.r.t. the reliability index [24], evidence codes were joined into top-level groups as suggested by [25]. As the compartment dataset consists of missing values corresponding to a single gene, duplicated entries were removed based on the most evident results, i.e., the higher priority was assigned to the entries verified by experiments and curated information. Subsequently, the subgroups were merged into the top-level group (e.g., “Organelle” and “Organelle part” into “Organelle”) in order to reduce the complexity of high-dimensional data. Most commonly-validated compartments were selected among duplicates. Finally, compartment names were updated by the corresponding GO term indices (”Membrane” localization into “GO:0016020” GO term into “16020” GO encoded) in order to preserve associations between the categories.

#### Imputation of missing values in generated feature matrix

Since some of the features (Table S2) consist of missing values, these values were imputed using a k-Nearest Neighbors (k-NN) approach. The exact behavior of the k-NN imputation model depends on the chosen distance metric. Here, given a sample containing a missing value indicator (NaN) for a feature, the value is being replaced by the average feature value calculated for *K* = 100 samples that are closest to the considered one, according to the Euclidean distance.

#### Calculation the averaged gene expression difference between experiments

In order to define a quantitative metric for comparing sequencing experiments, the aggregated per-sample gene expression difference was measured and used as a dependent variable for the regression analysis. Log-transformation of gene expression values was done to decrease variability within the sequencing data and make it conform more closely to the normal distribution. Gene expression data was represented as an *M* ^|*G*|*x*|*S*|^ matrix of genes vs. samples for S∈A∈B, where A and B are pseudo- and bulk RNA-Seq, respectively, and |*S*_*A*_| = |*S*_*B*_| = *N*. Then, the slope and intercept of the regression line can be estimated as shown in Equations 1-2, so that the sum of squared errors (SSE) is minimized (Equation 3). Then, the difference is measured as the observable error from the estimated coefficients of OLS.

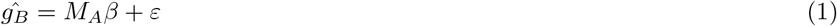

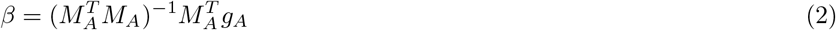

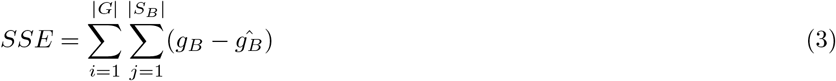

#### Assessment of feature importance using model-based feature selection

To choose the most suitable model w.r.t. feature selection, we trained and evaluated different machine learning algorithms in 5-fold cross validation, as well as benchmarked them against the baseline estimator (linear and logistic regression models for regression and classification tasks, respectively) in order to access the goodness of predictions. Random forest is the one of embedded approaches that was employed to rank features w.r.t. their influence on changes in the quantitative difference and leverage performance of decision trees, while mitigating its tendency to overfitting. For the regression task, standard deviation served as a measure of impurity, those decrease has been defining hierarchical split of the training samples [26]. Afterwards, we estimated importance of a feature proportionally to its closeness to the tree’s root node, which is defined through the information gain [27]. Finally, averaging by trees in the forest produces the joint estimation of the feature importance. In order to rank features w.r.t. their relevance for dropouts (classification task), we applied an MLP model [28]. Assessment of feature importance scores using this model was done through the accumulation of absolute values of gradient of the loss w.r.t. input during the model training [29].

Optimization of hyper-parameters of regression and classification models by the 5-fold CV grid-search over all possible combinations in the parameter space as the following: 1) Split the data into 4 training and 1 test folds. 2) Perform feature pre-processing to make values follow the normality assumption. 3) Train the model on the training folds and evaluate on the test fold. 4) Choose hyper-parameters corresponding to the best model. Hyper-parameter search for the random forest model was done for the number of estimators (from 32 to 1024), their maximum depth (4-32), minimum number of samples required to split an internal node (2-16) and the minimum number of samples in a leaf node (1-8). Optimization of hyper-xparameters for the MLP model was done for the number of neurons in hidden layers (32-1024), learning rate (0.001-0.1), and L2-regularization term (*α*; 0.0001-0.1). The MLP’s output was mapped non-linearly using logistic activation function, and its weights were adjusted during the training using Adam optimizer [30].

#### Identification of the core relevant features using recursive feature elimination

To identify an optimal subset of the features relevant for the gene expression difference between matched RNA-Seq experiments, we applied recursive feature elimination (RFE) [31], which performs an iterative elimination of least scored features w.r.t. to an external estimator (e.g., random forest). In contrast to the full search through all possible feature combinations, the RFE-based selection solves the task in a linear time, thus providing a valuable speed-up.

The optimal number of features is determined by computing index of the first stationary point of the objective function based on the RFE-provided objective values 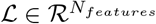 using first- (Δ) and second-order (Δ^2^) differences (Equation 4).

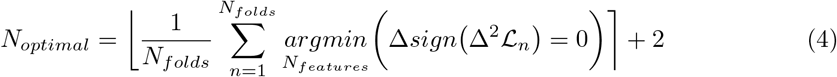

#### Statistical analysis

The correlation analysis is based on Spearman’s rank and Pearson correlation coefficients.

## Supplementary Information

### Additional Files

**Additional file 1**. The Supplemental Material includes Figures S1 to S2 and Tables S1 to S2.

### Availability of data and materials

The raw data of matched RNA-Seq experiments supporting the findings of this study are available at the European Nucleotide Archive (ENA) under accession number PRJEB43819. The source code of FAVSeq applied in this study is available on the GitHub repository (https://github.com/slipnitskaya/FAVSeq).

### Authors’ contributions

H.K. and K.B. devised the project, encouraged investigation of matched RNA-Seq data, suggested research challenges to be solved. S.L. designed and implemented the computational framework, performed data mining and ML-assisted analysis, wrote the manuscript with input from all authors. S.Le. provided expert advice and thoughtful suggestions to the manuscript. K.B. considerably contributed to the manuscript structure, carried out preliminary exploration of raw scRNA-Seq data and TPM normalization of unprocessed reads. Y.S. performed the pre-processing and enhancements of upstream RNA-Seq pipelines of raw sequencing reads and provided protocol assistance. All authors provided critical feedback and helped shape the research, analysis and manuscript.

