## Additional file 1 for "Machine learning-assisted identification of factors contributing to the technical variability between bulk and single-cell RNA-seq experiments"

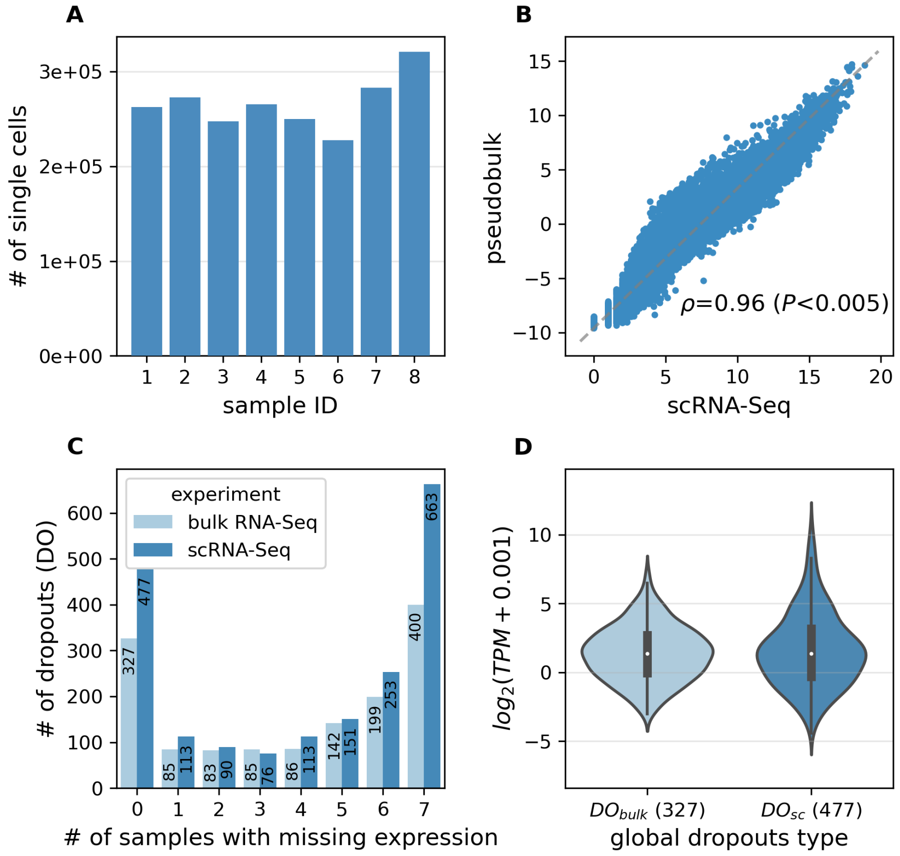


**Figure S1** Distributional statistics calculated for the matched single-cell and bulk RNA-Seq data. A. Number of cells measured in scRNA-Seq experiment across individual samples. B. Comparison of averaged gene expression measurements between scRNA-Seq counts and aggregated pseudobulk data. Log-transformed values are shown. C. Sparsity rate of dropouts calculated across different proportions of samples for each experiment. The middle-bar-number represents the amount of genes with missing expression observed in only one of the experiments, while being expressed in at least one sample in the matched one (sample-wise dropouts). D. Mean expression levels of genes that are global dropouts or represent missing expression in all samples in only one of the matched RNA-Seq experiments, while being detected across all samples by another technology (bars 0 in Figure S1C). Gene expression measurements in the matched experiment are shown.

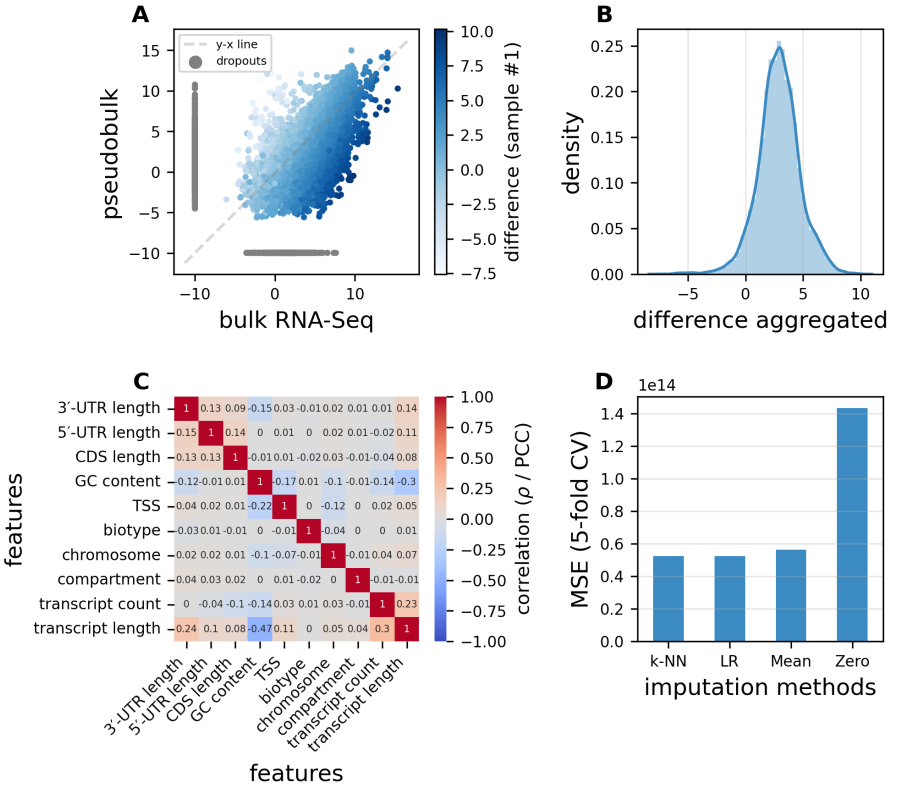


**Figure S2** Exploratory analysis of generated features and the target difference used in the FAVSeq pipeline. A. Gene expression difference—quantitative and dropouts-related—between the bulk and aggregated scRNA-Seq datasets measured in an individual sample for the most confident 13,427 genes detected in either of the matched experiments. Blue color indicates the most confident common genes detected in both matched experiments and gray indicates global dropouts (12,623 and 804 genes, respectively). Log-transformed values are shown. B. Distribution of the quantitative difference aggregated across 8 samples and calculated for the most confident common genes expressed in both RNA-Seq experiments. C. Cross-correlation analysis of gene-specific features filtered w.r.t. missing values (sparsity 0% in Table S2) and generated for the most confident 5,134 genes detected in either of RNA-Seq experiments. Color bar represents the distribution of correlation coefficients (Spearman rank and Pearson in lower left and upper right triangles, respectively), with grey color indicating values close to 0 (weak correlation). D. Performance of imputation approaches (k-Nearest Neighbors as k-NN, linear regression as LR, mean- and zero-based methods) done based on 5-fold CV in the feature reconstruction task and performed for all features with 20% of NaNs introduced manually in the training subset. Y-axis represents MSE that serves as an average error measure (lower is better) for the evaluated imputation methods.

**Table S1** Comparison of experimental protocols of matched RNA sequencing experiments

| Processing step | bulk RNA-Seq | scRNA-Seq |
|---|---|---|
| **Library type** | single cell library | single cell library |
| **Library preparation** | NEBNext Ultra II RNA 2x75bp | Single Cell 3' v2 |
| **Sequencing type** | dual-indexed paired-end | single-indexed paired-end |
| **Sequencing platform** | Illumina HiSeq 4000 | Illumina HiSeq 4000 |
| **Aligner** | STAR (v2.7.3a) | STARSolo (v2.7.3a) |

**Table S2** Sparsity of features containing missing values for different genes subsets

| Feature name | Genes with missing values (#) | |
|---|---|---|
| | **Sparsity 17.5%** | **Sparsity 0%** |
| 5'-UTR length | 4 266 | – |
| 3'-UTR length | 3 830 | – |
| transcript counts | 3 806 | – |
| GC content | 3 806 | – |
| TSS | 3 806 | – |
| compartment | 3 631 | – |
| CDS length | 385 | – |
| **Genes total (#)** | **13 427** | **5 134** |
| **Dropouts (#)<br>(minority class, %)** | **804<br>(6.0 %)** | **36<br>(0.7 %)** |

Sparsity: ratio of missing values (NaNs) in the feature space. 0 % indicates cleaned up data (no NaNs).
